# Molecular species selectivity of lipid transport creates a mitochondrial sink for di-unsaturated phospholipids

**DOI:** 10.1101/2020.06.08.140129

**Authors:** Mike F. Renne, Xue Bao, Margriet W.J. Hokken, Adolf S. Bierhuizen, Martin Hermansson, Tom A. Ewing, Xiao Ma, Ruud C. Cox, Jos F. Brouwers, Cedric H. De Smet, Christer S. Ejsing, Anton I.P.M. de Kroon

## Abstract

Mitochondria depend on the import of phospholipid precursors for the biosynthesis of the non-bilayer lipids phosphatidylethanolamine (PE) and cardiolipin required for proper function, yet the mechanism of lipid import remains elusive. Pulse labeling yeast with stable isotope-labeled serine followed by mass spectrometry analysis revealed that mitochondria preferentially import di-unsaturated phosphatidylserine (PS) for conversion to PE by the mitochondrial PS decarboxylase Psd1p. Several protein complexes tethering mitochondria to the endomembrane system have been implicated in lipid transport in yeast, including the endoplasmic reticulum (ER)-mitochondrial encounter structure (ERMES), ER-mitochondria complex (EMC) and the vacuole and mitochondria patch (vCLAMP). By limiting the availability of unsaturated phospholipids through overexpression of the glycerol-3-phosphate acyltransferase Sct1p, conditions were created to investigate the mechanism of lipid transfer and the contribution of the tethering complexes *in vivo*. Under these conditions, inactivation of ERMES components or the vCLAMP component Vps39p exacerbated the lipid phenotype, indicating that ERMES and Vps39 contribute to the mitochondrial sink for unsaturated acyl chains by mediating transfer of di-unsaturated phospholipids. The results support the concept that intermembrane lipid flow is rate-limited by molecular species-dependent lipid efflux from the donor membrane and driven by the lipid species’ concentration gradient between donor and acceptor membrane.

## Introduction

Mitochondria are essential cell organelles required for a plethora of functions, including energy production and lipid metabolism. Whereas most enzymes catalyzing the synthesis of bulk membrane lipids are localized to the endoplasmic reticulum (ER), evolution left PS decarboxylase (PSD) converting phosphatidylserine (PS) to phosphatidylethanolamine (PE), in mitochondria (Fig 1; [1]). Accordingly, PE is enriched in mitochondrial membranes compared to other organelles [2]. Mitochondria also harbor the enzymes involved in the biosynthesis of the mitochondrial signature lipid cardiolipin (CL) [3]. For the synthesis of PE and CL, mitochondria depend on import of the respective lipid precursors PS and phosphatidic acid (PA) from the ER [4].

**Figure 1.**
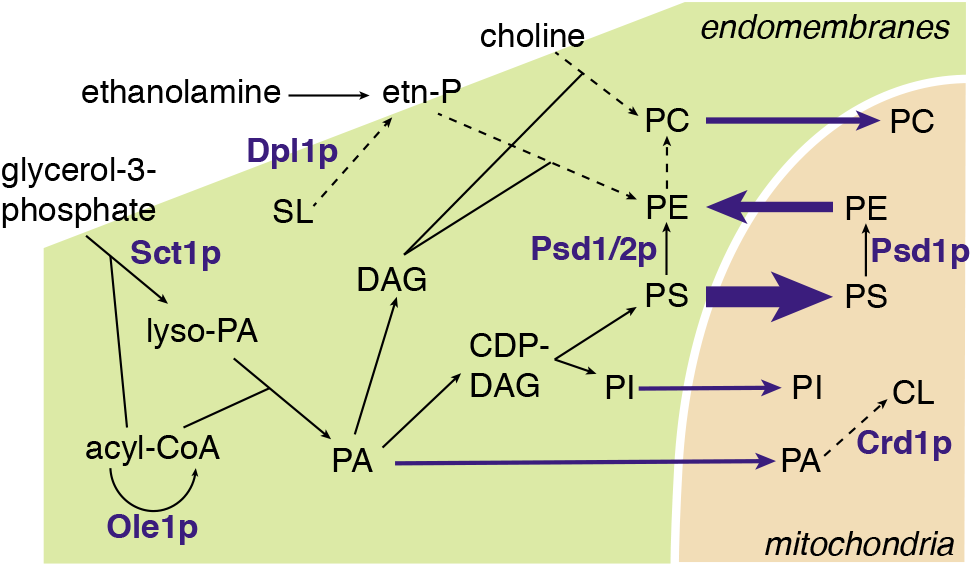
Overview of yeast phospholipid biosynthesis in endomembrane system and mitochondria, highlighting the enzymes most relevant to this study. Dashed arrows indicate multistep reactions; blue arrows indicate lipid flow into and from mitochondria. CL, cardiolipin; DAG, diacylglycerol; etn-P, phosphoethanolamine; PA, phosphatidic acid; PC, phosphatidylcholine; PE, phosphatidylethanolamine; PI, phosphatidylinositol; PS, phosphatidylserine; SL, sphingolipids.

Both PE and CL are membrane lipids with non-bilayer propensity that confer negative intrinsic curvature to membranes, the extent increasing with higher content of unsaturated acyl chains [5]. Loss of mitochondrial synthesis of PE and CL is synthetically lethal in yeast [6], thus a minimum level of non-bilayer preferring lipids is considered essential for mitochondrial function [7]. The enrichment of PE and CL in unsaturated acyl chains [8,9] further supports this notion. Local mitochondrial biosynthesis of PE is required for optimal respiration in yeast [10–12], and for potentiating respiratory capacity of skeletal muscles [13], highlighting the importance of the mitochondrial localization of PS decarboxylase. Defective PS transfer to mitochondria inhibiting mitochondrial PE synthesis, leads to liver disease [14].

Mitochondrial PSD enzymes are evolutionarily conserved [15]. In yeast, the mitochondrial PSD Psd1p catalyzes up to 90% of PS to PE conversion [16,17], with the remaining activity provided by Psd2p that is localized in endosomes [18]. Part of the PE produced in mitochondria relocates to the ER to be methylated to phosphatidylcholine (PC). As a consequence, yeast cells rely heavily on lipid transport between ER and mitochondria for supply of the major membrane lipids PE and PC, particularly in the absence of ethanolamine and choline, substrates of the CDP-ethanolamine and CDP-choline routes, respectively, that produce PE and PC in the ER (Fig 1; [19]). A twist to the longstanding concept of PSD activity residing exclusively outside the ER, came with the recent finding of a dual localization of Psd1p to mitochondria and ER, the distribution depending on metabolic demand [11].

To facilitate lipid transport, mitochondria maintain organelle contact sites (OCS) with an ER subfraction called mitochondria associated membranes (MAM) that is enriched in lipid biosynthetic enzymes [20], and with the vacuole [4]. The protein complexes that tether mitochondria to the ER, *i.e*. the ER-mitochondrial encounter structure (ERMES; [21,22]) and the ER membrane protein complex (EMC; [23]), as well as those that tether the ER to the vacuole, *i.e*. Vps13p-Mcp1p, and the vacuole and mitochondria patch (vCLAMP; [24–28]) have been implicated in lipid transport, of PS in particular. Recently, both a heterodimer of the ERMES subunits Mmm1p and Mdm12p [29,30] and the vacuole contact site protein Vps13p [31] were shown to mediate lipid transfer *in vitro*. However, the contribution of each tethering complex to lipid transport, and the molecular details of the process remain elusive.

The present study addresses the role of mitochondrial lipid import in establishing cellular lipid composition. Using a dynamic lipidomics approach we show that PS-to-PE conversion mainly produces di-unsaturated PE molecular species. This preference depends on the mitochondrial localization of Psd1p, indicating that lipid transport is molecular species selective, and explaining why mitochondria serve as a sink for unsaturated acyl chains. Next, the mechanism of mitochondrial lipid import including the role of OCS is interrogated by manipulating the cellular level of lipid unsaturation through overexpression of *SCT1. SCT1* encodes a glycerol-3-phosphate acyltransferase (Figure 1) that prefers C16:0-CoA as substrate [32]. Under conditions of *SCT1* overexpression, C16:0 acyl chains are sequestered in glycerolipids and protected from desaturation by the single essential Δ9-desaturase Ole1p, thus increasing the cellular content of saturated (SFA) at the expense of unsaturated acyl chains (UFA) [33]. By limiting the availability of di-unsaturated phospholipids, the overexpression of *SCT1* reveals the roles of the above tethering complexes in intermembrane lipid transfer *in vivo*, sheds light on the molecular mechanism of mitochondrial lipid import, and underscores the role of mitochondria in cellular membrane lipid homeostasis.

## Results

### PS decarboxylation is molecular species selective

To investigate PS-to-PE conversion by Psd1p at the level of individual lipid molecules, the incorporation of stable isotope-labeled ^13^C3^15^N-serine into PS, and the subsequent decarboxylation to ^13^C_2_^15^N-PE after a 20 min pulse was monitored by high resolution mass spectrometry in an established set of PSD mutants [11]. In the mutants, Psd1p was either inactivated or left as the only source of cellular PE by deletion of *PSD2* and *DPL1*, the latter encoding a dihydrosphingosine lyase that generates phosphoethanolamine [34], an intermediate in the CDP-ethanolamine route (Figure 1). Strains were cultured in synthetic defined galactose medium (SGal) devoid of choline and ethanolamine to confer maximal dependence on PS decarboxylation for *de novo* synthesis of phospholipids. The non-preferred carbon source galactose chosen to enable expression from the *GAL* promoter in the forthcoming, rendered cells partly dependent on mitochondria for energy supply [35,36].

The growth phenotype on SGal of the mutants compared to WT was similar to that on synthetic defined glucose medium (SD) (Figure S1A) [11], with *psd1Δ* and the mutant with Psd1 confined to mitochondria (*psd1Δpsd2Δdpl1Δ* +Psd1(Mt)) showing delayed growth.

Analysis of stable isotope labeled PS (*PS) and PE (*PE) abundance after the 20 min pulse showed that PE synthesis is decreased by ± 66% in *psd1Δ* compared to WT, whereas in *psd2Δdpl1Δ* in which Psd1p activity is the only source of PE, PS-to-PE conversion was similar to WT (Figure 2A). In WT cells, newly synthesized PE was enriched in di-unsaturated species compared to newly synthesized PS (Figure 2B). In *psd1Δ*, the preferential decarboxylation of *PS 34:2 was retained, whereas the enrichment of *PE 32:2 was lost and compensated for by a rise in the proportion of *PE 34:1. The enrichment of di-unsaturated *PE in *psd2Δdpl1Δ* was similar to WT (Figure 2B), in agreement with Psd1p being the main PS decarboxylase [10,16,37]. In line with these observations, steady state PE and PC in WT and *psd2Δdpl1Δ* were enriched in di-unsaturated species compared to the steady state PS, mainly at the expense of 34:1. This enrichment was largely lost in *psd1Δ* (Figure S1B). Taken together, the data show that preferential decarboxylation of di-unsaturated PS species by Psd1p plays a major role in establishing the PE and PC molecular species profiles under the culture conditions applied.

**Figure 2.**
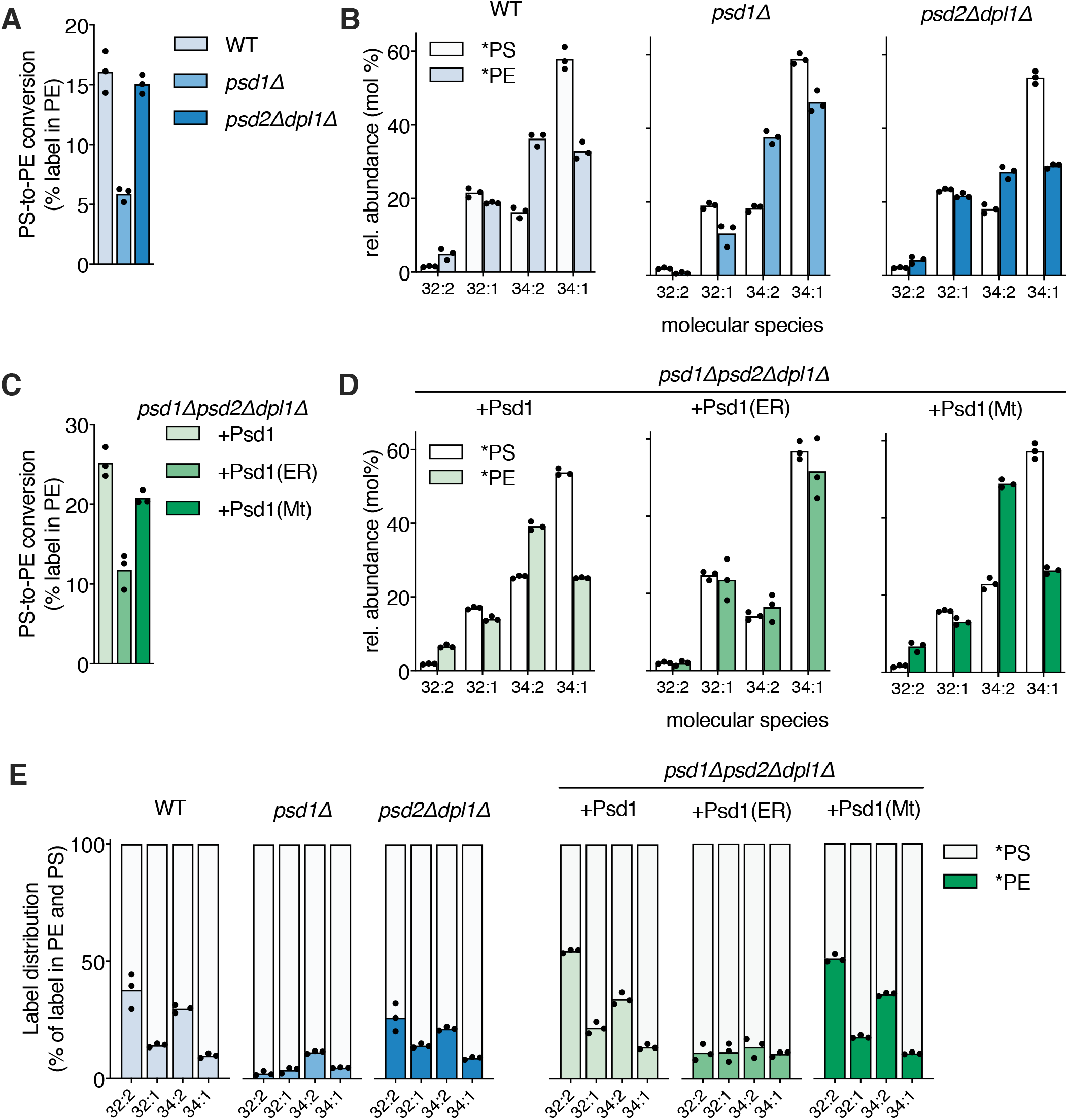
Molecular species selectivity of mitochondrial PS decarboxylation is due to species-selective transport rather than Psd1p substrate specificity. (A) PS decarboxylase activity in wild-type (WT, W303) and the indicated mutant strains expressed as percentage of the lipid-incorporated ^13^C^15^N-label recovered in PE, and (B) molecular species composition of ^13^C_3_^15^N-labeled PS (*PS) and ^13^C_2_^15^N-labeled PE (*PE) after 20 min incubation with ^13^C_3_^15^N-serine. (C) and (D) as (A) and (B) for the *psd1Δpsd2Δdpl1Δ* mutant expressing Psd1, Psd1(ER) or Psd1(Mt). (E) Molecular signatures of PSD activity represented by the label distribution per molecular species between *PS and *PE as percentages of the total amount of label incorporated in the molecular species indicated. Exponentially growing cells were pulsed for 20 min with ^13^C_3_^15^N-serine at 30°C prior to lipid extraction and lipid analysis by shotgun lipidomics. Molecular species (sum of carbon atoms in the acyl chains : sum of double bonds in the acyl chains) representing at least 1% of both labeled PS and labeled PE are shown in panels B, D (as mol% of total), and E. Compared to *PS and *PE, the amount of ^13^C_2_^15^N -PC synthesized was negligible (cf. [24]), and therefore not included in the analysis. Data are presented as the mean of 3 biological replicates, with the individual values indicated. Underlying data for this figure can be found in Data S1.

### Molecular species selectivity of mitochondrial PS decarboxylation is due to species-selective transport rather than Psd1p substrate specificity

The observed molecular species-selective conversion of PS by Psd1p could originate from substrate specificity, or from molecular species-dependent lipid transport as proposed previously [38,39]. To distinguish between these possibilities, we investigated the molecular species selectivity of PS-to-PE conversion in *psd1Δpsd2Δdpl1Δ* cells expressing chimeric Psd1p constructs that localize exclusively to the ER (Sec66-Psd1p chimera; +Psd1(ER)) or to mitochondria (Mic60-Psd1p chimera; +Psd1(Mt)) in comparison to WT Psd1p (+Psd1) [11]. Stable isotope labeling showed that PS-to-PE conversion proceeded at a higher rate in *psd1Δpsd2Δdpl1Δ* cells expressing Psd1 and Psd1(Mt) as compared to cells expressing Psd1(ER) (Figure 2C), as was reported previously [11]. The *psd1Δpsd2Δdpl1Δ* strains expressing Psd1 and Psd1(Mt) shared similar species profiles of newly synthesized PS and PE with a preference for the conversion of *PS 32:2 and *PS 34:2 to *PE (Figure 2D), as was observed in WT and *psd2Δdpl1Δ* (Figure 2B). Strikingly, in the *psd1Δpsd2Δdpl1Δ* strain expressing Psd1(ER), the *PS molecular species are converted to *PE with similar efficiencies (Figure 2D), demonstrating that the molecular species selectivity of PS decarboxylation depends on the mitochondrial localization of Psd1p. Moreover, Psd1p did not show any molecular species preference in decarboxylating endogenous and exogenously added PS molecular species in detergent-solubilized mitochondria (Figure S2), supporting Psd1p’s lack of intrinsic substrate preference.

The distribution of the label incorporated per molecular species between PS and PE provided the molecular signatures of PSD in the strains tested, and showed strong resemblance between the strains in which Psd1 has a mitochondrial localization (Figure 2E). The similarity of the signatures of *psd1Δpsd2Δdpl1Δ* cells expressing WT Psd1 and Psd1(Mt) argues that the contribution of the ER-localized Psd1p to PSD activity in the former, although essential for optimal growth (Figure S1A) [11], is quantitatively secondary. The differences in PS conversion between *psd1Δpsd2Δdpl1Δ* cells expressing Psd1 or Psd1(Mt) on the one hand versus *psd1Δpsd2Δdpl1Δ* expressing Psd1(ER) on the other, is also apparent from the steady state molecular species profiles with an enrichment of mono-unsaturated (32:1 and 34:1) PS, PE and PC in the latter (Figure S1C).

We conclude that the molecular species-selectivity of PS to PE conversion in yeast mitochondria is due to preferential import of di-unsaturated PS molecular species in agreement with previous findings in mammalian cells [39].

### Inactivation of mitochondrial lipid biosynthetic enzymes enhances the accumulation of saturated acyl chains under conditions of *SCT1* overexpression

To address the molecular mechanism of mitochondrial lipid import, we limited the cellular content of unsaturated acyl chains in wild type and mutants impaired in mitochondrial lipid biosynthesis and import, available in the BY4741 background. Overexpression of *SCT1* was previously shown to increase the cellular content of saturated acyl chains (SFA), C16:0 in particular [33]. Episomal overexpression of *SCT1* from a *GAL1* promoter accordingly increased mono- and di-saturated molecular species at the expense of di-unsaturated species in PS, PE, and PC in the BY4741 wild type strain (Figure S3). Of note, compared to W303 (Figure S1C), BY4741 (derived from FY1679) was enriched in C32 molecular species, which corresponds to the different acyl chain compositions of the two strain backgrounds [40].

Pulse labeling BY4741 with stable isotope-labeled ^2^H_3_-serine for 20 min revealed that, irrespective of *SCT1* overexpression, di-unsaturated PS species were more readily converted to PE than mono-saturated molecular species, similar as in W303 (Figure 3A and 2B), with superimposed a preference for decarboxylation of C32-over C34-PS species (Figure 3A). However, in the *SCT1*-overexpressing cells the proportion of newly synthesized di-unsaturated PE was decreased by 15% compared to the empty vector control.

**Figure 3.**
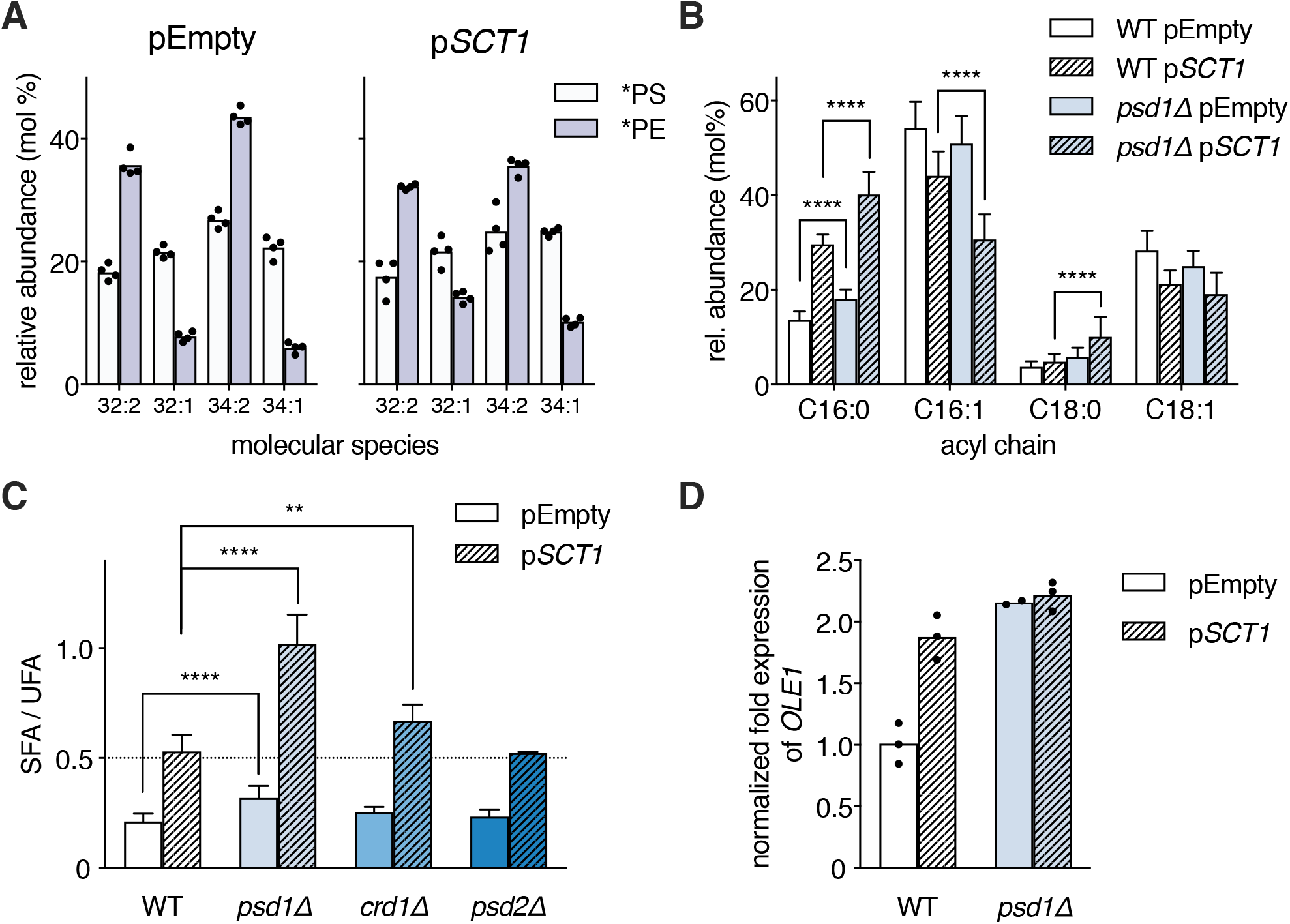
Loss of mitochondrial lipid biosynthetic enzymes enhances the accumulation of saturated acyl chains under conditions of *SCT1* overexpression. (A) The molecular species-selectivity of PS-to-PE conversion is recapitulated in wild type BY4741. WT cells overexpressing *SCT1 vs*. control were pulsed for 20 min with ^2^H_3_-serine prior to lipid extraction and ESI-MS/MS analysis of ^2^H_3_-PS (*PS) and ^2^H_3_-PE (*PE). Data is presented as mean of 4 biological replicates with the individual values indicated. Underlying data for this figure can be found in Data S2. (B) Acyl chain composition of WT and *psd1Δ* cells overexpressing *SCT1* (p*SCT1*) *vs*. empty vector control (pEmpty). Data is presented as mean ± SD (n>10). **** p < 0.0001, multiple t-test. (C) Ratio of the proportions of saturated over unsaturated acyl chains (SFA/UFA) of WT, *psd1Δ, crd1Δ* and *psd2Δ* overexpressing *SCT1 vs*. empty vector control. Data is presented as mean ± SD (n≥3). ** p<0.01, **** p < 0.0001, unpaired two-tailed t-test. (D) *OLE1* transcript levels in WT and *psd1Δ* transformed with p*SCT1 vs*. pEmpty as determined by RT-qPCR. Data were normalized to endogenous *ACT1* mRNA, and to the average of WT pEmpty (n=3).

Supplementation of the SGal medium with 0.05% glucose was required to allow proper growth of mutants disturbed in mitochondrial membrane biogenesis (Figure S4A,B). This modification of the culture medium had only minor effect on the rise in the proportion of SFA induced by overexpression of *SCT1* (Figure S4C). Importantly, interference with mitochondrial lipid biosynthesis by deleting *PSD1* aggravated the phenotype conferred by *SCT1* overexpression, as the proportions of C16:0 and C18:0 acyl chains were further increased at the expense of C16:1 and C18:1 (Figure 3B, Table S1), corresponding to a two-fold rise of the ratio of the proportions of saturated over unsaturated acyl chains (SFA/UFA) compared to WT (Figure 3C). Accordingly, the *psd1Δ* mutant overexpressing *SCT1* had a stronger growth phenotype than WT (Figure S4B). A much smaller, yet significant rise in SFA/UFA was observed in *psd1Δ* transformed with the empty vector compared to WT (Figure 3B,C), reflecting an increase in lipid saturation independent of *SCT1* overexpression, in agreement with the rise in 34:1 level in the PE and PC profiles of the *psd1D* mutant in the W303 background (Figure S1B). Enhancement of the accumulation of SFA induced by *SCT1* overexpression also occurred in *crd1Δ* cells that lack cardiolipin synthase, whereas it was absent in *psd2Δ* (Figure 3C), showing that the effect is not general for mutants in lipid metabolism.

The enhancement of the *SCT1* overexpression phenotype in *psd1Δ* and *crd1Δ* reflected increased incorporation of saturated acyl chains into lipids by Sct1p, outcompeting acyl-CoA desaturation by Ole1p [33]. Ole1p is known to be regulated at the level of transcription in response to changes in membrane physical properties [41,42]. Analysis by RT-qPCR showed that the increase in SFA levels in *psd1Δ* under *SCT1* overexpression was not due to a reduction in *OLE1* expression (Figure 3D). Instead, the *OLE1* transcript level was increased under conditions of *SCT1* overexpression and/or deletion of *PSD1*, probably in response to increased acyl chain saturation [43]. We propose that feed-back inhibition resulting from the diminished draw on the pool of unsaturated lipids and acyl-CoA’s caused by inactivation of Psd1p (or Crd1p), tipped the balance toward incorporation of saturated acyl chains into glycerolipids by Sct1p at the expense of desaturation by Ole1p. Conversely, the levels of newly synthesized di-unsaturated PS and PE went up with increasing mitochondrial Psd1 activity (Figure S5), indicating that increased draw enhanced the level of desaturation.

Taken together, the results define mitochondria as a sink for unsaturated acyl chains, in agreement with the above preferential conversion of di-unsaturated PS species to PE and the established enrichment of unsaturated acyl chains in mitochondrial membranes [2].

### ERMES and vCLAMP mutants overexpressing *SCT1* accumulate saturated acyl chains and exhibit a growth defect

If ERMES, EMC and vCLAMP contribute to mitochondrial phospholipid import, mutants lacking components of these tethering complexes are predicted to exhibit a reduction of the mitochondrial UFA sink. ERMES consists of Mmm1p, Mdm10p, Mdm12p, Mdm34p and a substoichiometric regulatory Miro GTPase Gem1p [21,22]. Under conditions of *SCT1* overexpression the ERMES mutants, *mmm1Δ, mdm10Δ, mdm12Δ, mdm34Δ* and *gem1Δ*, showed an increase of at least 50% in SFA/UFA ratio compared to WT. Accordingly, growth of ERMES mutants overexpressing *SCT1* was strongly retarded compared to WT and empty vector control (Figure 4B). In contrast, growth of single-, double- (not shown), and triple mutants of genes encoding subunits of the EMC tethering ER to mitochondria [23], was not affected by *SCT1* overexpression, while that of the quadruple mutant tested was slightly impaired (Figure 4B). Since the *3xemcΔ* and *4xemcΔ* mutants overexpressing Sct1p (Figure S6) did not show increased accumulation of SFA (Figure 4A), this was probably not due to defective PS transfer.

**Figure 4.**
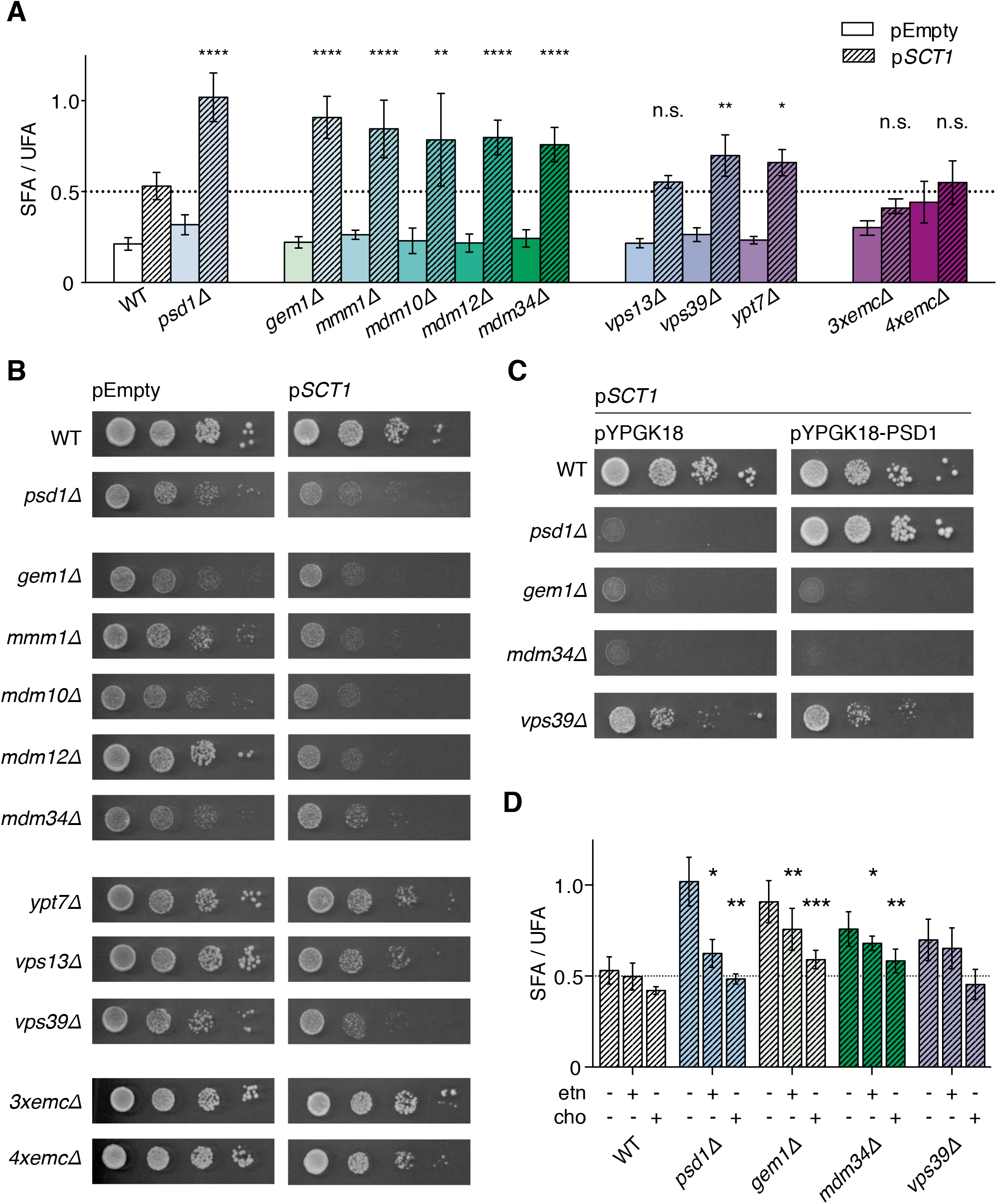
ERMES and vCLAMP mutants overexpressing *SCT1* accumulate saturated acyl chains and exhibit a growth defect that is partially rescued by exogenous ethanolamine and choline. (A) SFA/UFA ratios of WT and indicated mutant strains overexpressing *SCT1* (p*SCT1*) *vs*. control (pEmpty). Data is presented as mean ± SD (n?3). * p < 0.05, ** p < 0.01, *** p < 0.001, **** p < 0.0001, n.s. not significant, unpaired two-tailed t-test of the indicated bar compared to WT p*SCT1*. (B) Growth of WT and indicated mutant strains overexpressing *SCT1* (p*SCT1*) *vs*. pEmpty. Serial dilutions (10^−1^ – 10^−4^) were spotted on SGal + 0.05% glucose and incubated for 3 days at 30°C. (C) Growth of WT and indicated mutants overexpressing *SCT1* co-transformed with pYPGK18-*PSD1* expressing *PSD1* from the *PGK1* promoter or with the corresponding empty vector (pYPGK18). Serial dilutions (10^−1^ – 10^−4^) were spotted on SGal + 0.05% glucose plates and incubated for 4 days at 30°C. (D) SFA/UFA-ratio of WT and indicated mutant strains overexpressing *SCT1* cultured without (data taken from panel A) and with 1 mM ethanolamine (etn) or 1 mM choline (cho) as indicated. Data is presented as average ± SD (n≥3). Stars indicate p-values as above in panel A, determined by an unpaired t-test of the indicated bar compared to the corresponding condition of WT.

Mitochondria are tethered to the vacuole by vCLAMP consisting of Ypt7p, Vps39p and Tom40p, and by a second, independent OCS formed by Vps13p and Mcp1p [26]. The SFA/UFA ratio of *ypt7Δ* and *vps39Δ* exhibited an increase around 30%, while that of *vps13Δ* did not significantly differ from WT (Figure 4A, Table S1). vCLAMP mutant *vps39Δ* showed delayed growth upon overexpressing *SCT1*, whereas growth of *ypt7Δ* and *vps13Δ* was only slightly affected (Figure 4B).

To exclude that the phenotypes induced by overexpressing *SCT1* in ERMES- and vCLAMP-mutants resulted from indirect effects related to Psd1p activity or to the level of *SCT1* overexpression, the following controls were done. Western blot analysis showed that the expression level of Psd1p was not decreased, and the level of Sct1p overexpression not increased in the tether mutants *versus* the WT (Figure S6). Moreover, episomal expression of *PSD1* from a *PGK1*-promoter, while restoring growth of the *psd1Δ* strain, did not rescue the growth phenotype of the ERMES and vCLAMP mutants under conditions of *SCT1* overexpression (Fig 4C). Based on the results, the enhancement of *SCT1* overexpression-induced SFA-accumulation in the ERMES and vCLAMP mutants was attributed to impaired transport of di-unsaturated lipids.

### Rescue of ERMES mutants and *vps39Δ* overexpressing *SCT1* by exogenous ethanolamine and choline

Under the growth conditions used, *i.e*. culture media devoid of choline and ethanolamine, cells depend to a large extent on mitochondrial PSD activity, for net synthesis of PE and PC. Supplementation of ethanolamine and choline reduces the cellular requirement for mitochondrial lipid synthesis by diverting lipid flux into the CDP-ethanolamine- and CDP-choline branches of the Kennedy-pathway, respectively (Figure 1; [4]). Since both routes draw on the pool of unsaturated acyl-CoA’s via PA and diacylglycerol (DAG; [44]), choline and ethanolamine were expected to alleviate the feed-back inhibition of Ole1p. In *psd1Δ* overexpressing *SCT1*, choline restored the SFA/UFA ratio to WT level (Figure 4D). Ethanolamine did not completely reverse the phenotype (Figure 4D), indicating that the demand for acyl-CoA by flux through the CDP-ethanolamine pathway was insufficient to compensate for the reduced mitochondrial UFA sink. Compared to *psd1Δ*, ethanolamine only partially (if at all) decreased the accumulation of SFAs conferred by *SCT1*-overexpression in the ERMES mutants and *vps39Δ* (Figure 4D). Choline strongly suppressed SFA accumulation in all strains, with the SFA/UFA ratio in *vps39Δ* returning to WT and in *mdm34Δ* and *gem1Δ* staying slightly above (Figure 4D). The extents to which ethanolamine and choline reverse the slow growth phenotype of the *SCT1*-overexpressing mutants (Figure S7) paralleled the corresponding decreases in SFA/UFA ratio. The incomplete restoration of the SFA/UFA ratios by the supplements in the tether mutants suggests that ERMES facilitates PE and PC transfer and that Vps39 mediates transfer of PE.

## Discussion

To limit UFA availability and study its effect on lipid transfer to mitochondria, an artificial metabolic sink for SFA was introduced in yeast by overexpressing *SCT1*. The rise in SFA/UFA resulting from overexpressed Sct1p outcompeting Ole1p, serves as a sensitive read-out of interference with the mitochondrial UFA sink, as validated in *psd1D* and *crd1D* and demonstrated in ERMES mutants. The rise in SFA/UFA is most likely caused by feedback inhibition with the reduced demand for unsaturated lipids stimulating incorporation of SFA into lipids by overexpressed Sct1p at the expense of desaturation by Ole1p. Besides mass action, product inhibition of lipid biosynthetic enzymes as demonstrated for PS synthase [45], may contribute to this feed-back mechanism.

Stable isotope labeling combined with MS analysis revealed that shorter di-unsaturated PS molecular species are preferentially converted to PE by mitochondrial Psd1p (Figure 2B,3A). Since ER-localized Psd1p does not distinguish between PS molecular species (Figure 2D,E), the transfer of PS from ER to mitochondria confers the molecular species selectivity. The preferred decarboxylation of shorter and more unsaturated PS molecular species in mammalian cells was interpreted in terms of the rate of transfer to mitochondria being inversely proportional to the molecular hydrophobicity of the PS molecular species, which led to the hypothesis that the efflux from the ER membrane constitutes the rate-limiting step [39,46]. Taking advantage of *SCT1* overexpression to manipulate the acyl chain composition in yeast, this hypothesis was experimentally tested *in vivo*. The reduced synthesis of PE under conditions of *SCT1* overexpression [33] and the aggravation of the *SCT1* overexpression phenotype in ERMES and vCLAMP deletion mutants provide the first *in vivo* evidence for lipid efflux from the donor membrane being rate-limiting in intermembrane lipid transport.

Based on a comparison of SFA/UFA ratios (Figure 4A), using *psd1D* (with a strongly reduced mitochondrial sink for unsaturated phospholipids) as a reference, ERMES appears to be the main contributor to mitochondrial import of lipids, PS first and foremost [21], under the conditions used. EMC consists of multiple subunits and tethers to the TOM-complex in the mitochondrial outer membrane. Deletion of multiple EMC genes is synthetically lethal with ERMES mutations and slows down PS to PE conversion [23]. However, when judged by the criterion of SFA/UFA ratio under conditions of *SCT1* overexpression, EMC plays a minor role, if any, in PS import. Instead, the role of EMC in membrane protein biogenesis [47], may account for the reported aberrant lipid metabolism of EMC mutants.

Deletion of *VPS39* and *YPT7* was found to enhance the *SCT1* overexpression phenotype (Figure 4A), indicating that lipid transfer from the vacuole contributes to the mitochondrial sink for di-unsaturated phospholipids. Since Vps39 and Ypt7 are part of both vCLAMP and the HOPS complex that functions upstream supplying lipids to the vacuole by vesicular transport [26,48], the present data do not allow distinction between defective HOPS or vCLAMP exacerbating the *SCT1* overexpression phenotype. The lack of rescue by ethanolamine of *vps39D* overexpressing *SCT1* (Figure 4D) is in agreement with the recently identified third function of Vps39p, trafficking ethanolamine-derived PE from ER to mitochondria independent of vCLAMP or HOPS [49]. Inactivation of Vps13p did not affect the SFA/UFA ratio, consistent with its function in mitochondrial lipid import being redundant with that of ERMES and only becoming apparent in the absence of functional ERMES [26,27,50]. The defects in intermembrane transport caused by deficiency of ERMES or Vps39p under conditions of limiting UFA are not restricted to PS, but also pertain to PE and PC, evidenced by the SFA/UFA ratios not returning to WT level when ethanolamine or choline is supplemented (Figure 4D).

Model membrane studies have shown that the rate of desorption of a lipid from a bilayer is determined by the lipid’s molecular structure and the physical properties of the donor membrane (reviewed in [51]). Our results provide evidence for molecular species-selective lipid flow into mitochondria at OCS that is rate-limited by lipid release from the donor membrane, and facilitated in an additive manner by the lipid species’ concentration gradient between donor and acceptor membrane and by the presence of tethers that serve as passive conduits increasing the probability of transfer to the juxtaposed membrane (Figure 5). Since desorption favors di-unsaturated hydrophilic lipid species [51,52], their concentration gradient is of prime importance.

**Fig 5.**
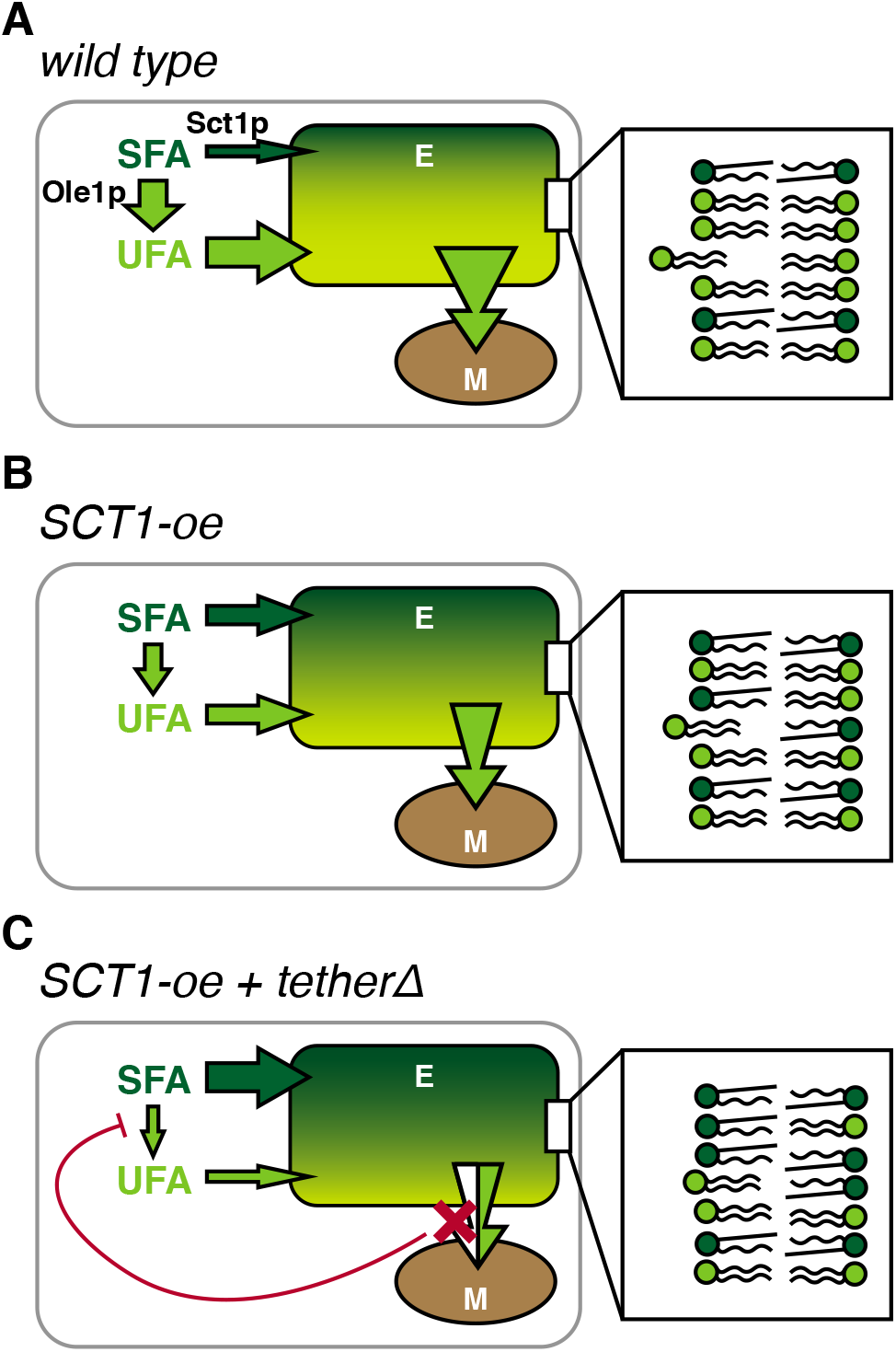
Model of the cross-talk between the SFA sink introduced by *SCT1* overexpression and the mitochondrial UFA sink in wild type yeast and mutants disturbed in ERMES or vCLAMP. Schematic representation of the relative contributions of Sct1p and Ole1p to the SFA (dark green)/UFA (light green) ratio, and its consequences for lipid flux from the endomembrane system (E) into mitochondria (M) in WT (A), WT overexpressing SCT1 (*SCT1*-oe) (B), and cells lacking a tether that overexpress *SCT1* (*SCT1*-oe *tetherΔ*) (C). In the latter, the flux of SFA into the pool of membrane lipids is further enhanced by the depicted feedback mechanism in response to partially obstructing (X) lipid entry into mitochondria. The insets show the tendency of a di-unsaturated lipid to efflux from the endomembrane system for each condition, depicted as the extent of displacement from the bilayer.

In case of PS, the gradient is maintained by the enrichment of PS synthesis in MAM [20], where local biosynthesis of PS at OCS has been shown to drive lipid transport [45], and by the efficient conversion of di-unsaturated PS to PE by mitochondrial Psd1p upon transfer. As a matter of fact, PS-to-PE conversion by mitochondrial Psd1p is more efficient than by Psd1 homogeneously distributed over the ER membrane (Figure 2A; [11]). Lowering the donor membrane’s UFA content by overexpression of *SCT1* reduces the steepness of the chemical gradient of di-unsaturated PS, as well as its rate of desorption by decreasing membrane fluidity (Figure 5A and B). As a result, yeast cells overexpressing *SCT1* produce reduced proportions of newly synthesized di-unsaturated PE and exhibit an overall reduced PE level [33]. Interference with the tethering complexes exacerbates the decrease in UFA content, most likely by causing feed-back inhibition of Ole1p activity that enables overexpressed Sct1p to sequester even more SFA in glycerolipids (Figure 5C).

ERMES subunits are equipped with synaptotagmin-like mitochondrial lipid-binding protein (SMP) domains that facilitate lipid transfer by providing hydrophobic binding pockets for the released lipid [30,53]. In agreement with lipid release from the donor membrane being rate-limiting for transfer, the endogenous lipids co-purifying with overexpressed Mdm12p are enriched in di-unsaturated, short 32:2 PC [53]. Concentration gradient-driven lipid transfer between membranes as proposed here is fully compatible with the hydrophobic channel structure of Vps13 [54].

An early link between cellular UFA content and mitochondrial function was the identification of the *mdm2* mutation that confers defective intracellular mitochondrial movement, as an *OLE1* allele [55]. In support of UFA availability limiting mitochondrial lipid import, the loss of fitness induced by deletion of *MDM34* was compensated by a point mutation in *MGA2* that confers two-fold upregulation of the *OLE1* transcript, and by supplementation with oleic acid [56]. Accordingly, the growth phenotype of ERMES mutants was alleviated by overexpression of *OLE1* [57].

Mitochondrial function requires the non-bilayer lipids PE and CL [7], and their non-bilayer propensity critically depends on UFA content [5]. The proposed concentration gradient-driven transfer of preferentially di-unsaturated PS and PA from ER to mitochondria by tether-facilitated diffusion, warrants optimal PE and CL molecular species composition (*cf*. [12]), explaining why PE and CL synthesis have been retained in mitochondria during evolution. The presence of multiple tethers connecting mitochondria to the endomembrane system, some with redundant functions in lipid transport, underscores the importance of this process.

The results presented have implications for our understanding of cellular membrane lipid homeostasis. The identification of mitochondria as a sink for phospholipids with unsaturated acyl chains accounts for the rise in unsaturated over saturated acyl chain content in yeast cells cultured on non-fermentable *versus* fermentable carbon source [2]. The molecular species-selective PS transfer to mitochondria explains the longstanding observation that newly synthesized PS is more readily converted to PE than pre-existing PS [20,58]. Moreover, it accounts for the higher SFA content of PS compared to PE and PC [8,59,60]. The continuous equilibration of newly synthesized PC and its di- and mono-methylated precursors between ER and mitochondria reported previously [61], further supports the notion of gradient-driven lipid transfer at membrane contact sites.

In conclusion, the dynamic manipulation of lipid acyl chain composition by overexpressing Sct1p has uncovered mechanistic details of mitochondrial lipid import, and identified the mitochondrial molecular species selectivity filter as a novel player in membrane lipid homeostasis.

## Materials and methods

### Strains, plasmids and culture conditions

Yeast strains used were derived from W303 and BY4741 and are listed in Table S2. Strains were maintained and pre-cultured in synthetic glucose medium (SD), containing per liter 6.7 g of yeast nitrogen base (Difco, B/D Bioscience), 20 g of glucose, 20 mg of adenine, 20 mg of arginine, 20 mg of histidine, 60 of mg leucine, 230 mg of lysine, 20 mg of methionine, 300 mg of threonine, 20 mg of tryptophan, and 20 mg uracil. Strains containing plasmids listed in Table S2, were cultured in drop-out media devoid of the appropriate components.

The *PSD1*-expression vector was constructed from pYES2-*PSD1*-HA by releasing the *PSD1-HA* fragment by double digestion and subsequent ligation into pYPGK18. Correct insertion was verified by restriction analysis and Psd1p expression was verified by complementation of the *psd1Δ* mutant.

Overnight cultures were diluted in SD and cultured to exponential growth. Cells were harvested, washed with water, and inoculated at OD_600_ < 0.05 in pre-warmed synthetic galactose medium (SGal), and cultured overnight. SGal contained per liter 6.7 g in-house mixed inositol- and choline-free YNB [62], 20 g of galactose (Acros Organics), 13.5 mg *myo*-inositol (final concentration 75 μM), and amino acids as above, with or without 0.5 g glucose as indicated. Strains were cultured at 30°C while shaking (200 rpm) and harvested during exponential growth (OD_600_ 0.3 – 0.8). Cell pellets were stored at −20°C until further processing.

Growth phenotypes were determined using a serial dilution spot assay. Cells were grown overnight in SD, diluted and cultured to exponential growth (OD_600_ 0.5 – 1.0). A culture volume corresponding to 0.5 OD_600_ units was harvested, washed with water, resuspended in water at OD_600_ of 1, and serially diluted in 10-fold increments to 10^−4^. 4 μL aliquots of each dilution were spotted on solid SGal medium and incubated at 30°C for the number of days indicated.

### ^13^C_3_^15^N-serine pulse labeling and lipid analysis by shotgun lipidomics

Yeast strains were cultured to early exponential growth (OD_600_ 0.4 – 0.6) in 100 mL SGal medium as above. ^13^C_3_^15^N-serine (Cambridge Isotope Laboratories) was added to a final concentration of 100 mg/L. After 20 min cellular metabolism cellular metabolism was quenched by the addition of perchloric acid (70%, Sigma Aldrich) to a final concentration of 0.66 M and rapid cooling in an ice-water bath. Cells were pelleted, washed 3x with 155 mM ammonium bicarbonate, snap frozen in liquid nitrogen and stored at −20°C until further processing. Lipids were extracted using a two-step extraction procedure [9] and analyzed by shotgun lipidomics [9,63]. Briefly, cells were dispersed at 5 OD_600_ units per mL in 155 mM ammonium formate and lysed using glass beads. Lipids were extracted from cell lysates corresponding to 0.4 OD_600_ units in 200 μL buffer that were spiked with a cocktail of lipid standards. 990 μL chloroform/methanol (17:1, v/v) was added followed by vigorous shaking for 120 min at 4°C. The lower organic phase was collected after centrifugation. The remaining aqueous phase was re-extracted with 990 μL chloroform/methanol (2:1, v/v) for 60 min, and the lower organic phase was collected. Lipid extracts were vacuum dried and dissolved in chloroform/methanol 2:1 (v/v). Lipid extracts were analyzed in negative and positive mode on an Orbitrap Fusion Tribrid mass spectrometer (Thermo Fischer Scientific) equipped with a Triversa Nanomate (Advion Biosciences) [64]. Lipidomics data analysis was performed using ALEX^123^ software [65,66].

### ^2^H_3_-serine pulse labeling and lipid analysis by tandem MS

Wild type yeast harbouring pYES2-*SCT1*-HH or pYES2-Empty was cultured to early exponential growth (OD_600_ 0.4 – 0.6) in 100 mL SGal -ura medium as above. At time zero deuterium-labeled, ^2^H_3_-serine (Cambridge Isotope Laboratories) was added to a final concentration of 100 mg/L. After 20 min cellular metabolism was quenched by addition of trichloroacetic acid to a final concentration of 5% (w/v) and rapid cooling on ice. Cells were pelleted, washed with water and stored at −20°C until further processing. Lipids were extracted as detailed below, dried under nitrogen and stored at −20°C. For ESI-MS/MS analysis, lipid extracts were directly injected and analyzed on an API4000 triple quadrupole instrument (Applied Biosystems). Detection of PS and ^2^H_3_-PS was in negative mode using neutral loss scans for 87 and 90 amu, respectively. PE and ^2^H_3_-PE were detected in positive ion mode by neutral loss scans for 141 and 144 amu, respectively. PC was detected in the positive mode by parent ion scanning for *m/z* 184. Data were converted to mzML format and analyzed using XCMS version 1.52.0 [67] running under R version 3.4.3. All data was isotope corrected as described [68].

### *In vitro* Psd1p assay

The *psd2Δ* strain was cultured in semi-synthetic lactate medium [69] to late exponential growth (OD_600_ 2.5), harvested and washed with 1 mM EDTA. Spheroplasts were prepared as described (Daum et al., 1982) using Zymolyase 100T (Seikagaku; 0.5 mg/g cells wet weight). The spheroplasts were resuspended in ice-cold D-buffer (0.6 M sorbitol, 10 mM Tris-HCl pH 7.4) containing protease inhibitors (complete protease inhibitor cocktail, Roche; 1 tablet per 50 mL), and lysed by 10 strokes in a Dounce homogenizer. Unbroken cells and nuclei were removed by two low-spin centrifugation steps (1400 g and 1500 g, respectively, 4 min, 4°C). The mitochondrial fraction was pelleted (9700 g, 10 min, 4°C), resuspended in assay buffer (0.6 M sorbitol, 1 mM EDTA, 50 mM Tris-HCl pH 7.4), and stored at −80°C. Protein concentration was determined using the BCA assay (Pierce) in the presence of 0.1% SDS with BSA as standard.

*In vitro* PS decarboxylation was analyzed at 0.5 mg/mL total mitochondrial protein in assay buffer also containing 0.1% (w/v) Triton X-100. Synthetic PS molecular species (Avanti Polar Lipids) were added at 50 μM from 10x stocks in assay buffer containing 0.1% Triton X-100. The reaction mixture was incubated at 30°C (with vigorous shaking) and 100 μL samples were drawn at t=0 and t=1 h and the reaction was stopped by adding these samples to chloroform/methanol (1:1) and rapid cooling on ice. Lipids were extracted, dried under nitrogen, and stored at −20°C.

Lipid extracts were analyzed for PS and PE by LC-MS. Chromatography was performed on a hydrophilic interaction liquid chromatography (HILIC) column (2.6 μm HILIC 100 Å, 50 x 4.6 mm, Phenomenex, Torrance, CA), by elution with a gradient from acetonitrile/acetone (9:1, v/v) to acetonitrile/water (7:3 v/v, containing 10 mM ammonium formate), both containing 0.1% (v/v) formic acid, at a flow rate of 1 mL/min. The column outlet of the LC was connected to a heated electrospray ionization (hESI) source of an Orbitrap Fusion Tribrid mass spectrometer (Thermo Fischer Scientific). Full spectra were collected from *m/z* 400 to 1150 at a resolution of 120,000 in negative mode. Data were converted to mzML format and analyzed as above [67,68].

### Lipid extraction

Cell pellets were resuspended in 600 μL water, transferred to cooled 2 mL Eppendorf tubes containing 0.5 mL of acid washed glass beads (Sigma), and bead-bashed for 5 min at RT (Qiagen TissueLyser II). Lipid extracts were prepared using a modified version of the Bligh and Dyer lipid extraction [70]. Briefly, the 600 μL cell lysate was transferred to a glass tube containing 1360 μL methanol and 620 μL chloroform, and 20 μL 1 M HCl was added. The mixture was vortexed briefly and kept on ice for 2 min. Next, 600 μL chloroform and 600 μL 0.1 M HCl were added, the mixture was vortexed briefly, kept on ice for 2 min and centrifuged for 4 min at 3000 g at 4°C to induce phase separation. The organic phase was collected. After adding 120 μL 1 M KCl to the water phase, it was washed with 600 μL chloroform, and the organic phases were pooled. The combined organic phase was washed with 0.1 M KCl, and after adding 120 μL isopropanol dried in a water bath (40°C) under N_2_-flow. Phospholipid concentrations were determined according to [71], after destruction in 70% perchloric acid for 1 h at 180°C.

Lipid extracts from the *in vitro* Psd1p assay samples were prepared as above using 6x smaller volumes.

### Analysis of acyl chain composition by GC-FID

Total lipid extracts corresponding to 200 nmol phospholipid phosphorus were transesterified by heating for 2.5 h at 70°C in 2.5 mL 2.5% (v/v) methanolic H_2_SO_4_. Fatty acid methyl esters (FAME) were extracted by addition of 2.5 mL hexane and 2.5 mL water. After collecting the organic phase, the aqueous phase was washed with hexane, the organic phases were pooled and washed with water until the pH of the water phase was neutral. 120 μL isopropanol was added and the sample was dried in a water bath (40°C) under N_2_-flow. The FAME were dissolved in 1 mL hexane, transferred to an Eppendorf tube and centrifuged for 10 min (14,000 g at 4°C) to remove any residual particles. Next, 900 μL of the supernatant was concentrated to a volume of 50 - 100 μL under N_2_-flow.

FAME were analyzed by GC-FID using splitless injection (2 μL injection volume, inlet temperature 230°C, 1 min splitless time with a split flow of 20 mL/min) on a Trace*GC Ultra* (Thermo Scientific) equipped with a biscyanopropyl-polysiloxane column (Restek), using N_2_ as carrier gas (1.3 mL/min, constant flow). After injection, the samples were concentrated on the column by keeping the temperature at 40°C for 1 min, after which the column was rapidly heated to 160°C (30°C/min). FAME were separated using a temperature gradient from 160°C to 220°C (4°C/min) and signal was detected using FID at 250°C (H_2_/air with N_2_ as make-up).

Integrated peaks were identified and calibrated using a commercially available FAME standard (63-B, Nu-Chek-Prep). Acyl chain compositions are presented as mol% of the four most abundant acyl chains (C16:0, C16:1, C18:0 and C18:1) that were recovered in all samples.

### Analysis of *OLE1* expression by RT-qPCR

A cell pellet corresponding to 20-25 OD_600_ units was lysed with glass beads and RNA was isolated from the cell lysate using the RNeasy Mini Kit and RNase-free DNase Set (both Qiagen) according to the manufacturer’s instructions. RNA quality and quantity were checked by agarose gel electrophoresis and spectrophotometry using a NanoDrop 2000 spectrophotometer (ThermoFischer Scientific), respectively. cDNA was synthesized from 1 μg RNA using Superscript III Reverse Transcriptase (Invitrogen) and oligo (dT)_12-18_ primers according to the manufacturer’s instructions. RT-qPCR was performed using TaqMan Universal PCR mastermix 2x (Applied Biosystems, ThermoFischer Scientific) supplemented with AmpErase UNG (Applied Biosystems, ThermoFischer Scientific), using commercially available TaqMan probes and primers for *OLE1* (Sc04122147_s1) and *ACT1* (Sc04120488_s1, ThermoFischer Scientific). PCR reactions were run on a ViiA 7 Real Time PCR machine (ThermoFischer Scientific) and data was analyzed using corresponding software. Ct values were averaged from two technical duplicates, and cDNA-devoid reactions were included as negative control. Gene expression was analyzed as 2^−ΔΔCt^ [72], and normalized to *ACT1* as housekeeping gene.

### Western Blot analysis

Proteins were extracted from cell pellets by NaOH treatment and boiling in Laemmli sample-buffer as described [73]. Aliquots were taken for acetone precipitation followed by solubilization in urea/guanidine and protein determination using the Bradford assay as described [74]. Subsequently, samples corresponding to equal amounts of protein were separated on a 10% SDS-PAGE gel and transferred to a nitrocellulose membrane by the wet tank blot method (BioRad, according to manufacturer’s instructions). Proteins were detected using primary antibodies against Sct1p-HA (anti-HA, Thermo Scientific), Psd1p [75], and Gpd1p (anti-GAPDH, Thermo Scientific), and IRdye-conjugated secondary antibodies (Li-Cor Biosciences, Lincoln, NE), and imaging on a Li-Cor Odyssey infrared imager.

### Statistical analysis

Acyl chain composition was analyzed using a multiple t test, analyzing each acyl chain percentage individually and using the False Discovery Rate approach (Q=5%). SFA/UFA ratios were analyzed by unpaired, parametric t-test. All statistical analyses were done using GraphPad Prism 6 software.

## Author contributions

Conceptualization: M.F.R., C.H.D.S., A.I.P.M.d.K.; Funding acquisition: M.F.R., X.B., C.S.E., A.I.P.M.d.K.; Investigation: M.F.R., X.B., M.W.J.H., A.S.B., M.H., T.A.E., X.M., R.C., J.F.B., C.H.D.S.; Methodology: M.F.R., X.B., J.F.B., C.S.E., A.I.P.M.d.K.; Supervision: M.F.R., A.I.P.M.d.K., Visualization: M.F.R., A.I.P.M.d.K., Writing-original draft: M.F.R., A.I.P.M.d.K.; Writing-review & editing: all authors.

## Acknowledgements

We are indebted to Dr. Jodi Nunnari for sharing the PSD mutants. We thank Dr. William Prinz, Dr. Benoît Kornmann, Dr. Steven Claypool, Dr. Takashi Tatsuta, Dr. Karin Athenstaedt and Dr. Günther Daum for kindly providing strains, plasmids and antibodies. This research was supported by the Division of Chemical Sciences in the Netherlands, with financial aid from The Netherlands Organization for Scientific Research (711-013-004, MFR), by the China Scholarship Council (grant no. 201204910146, XB), by the Barth Syndrome Foundation (AdK), by a Summer Fellowship from the Federation of European Biochemical Societies (MFR), by the VILLUM Foundation (VKR023439, C.S.E.), the VILLUM Center for Bioanalytical Sciences (VKR023179, C.S.E.) and the Lundbeckfonden (R54-A5858, C.S.E.).

## Supporting information

**Table S1. Acyl chain composition of WT and indicated mutants under *SCT1*-overexpression (p*SCT1*) *vs*. empty vector control (pEmpty)**

(DOCX)

**Table S2. Yeast strains and plasmids**

(DOCX)

**Data S1. Complete dataset of PS, PE and PC molecular species in W303 and derived mutants after 20 min labeling with ^13^C_3_^15^N-serine, averaged as mol% of total PS-PE-PC (± SD, n=3)**

(XLSX)

**Data S2. Complete dataset of PS, PE and PC molecular species in BY4741 p*SCT1 vs*. pEmpty, including ^2^H_3_-labeled PS and PE after 20 min labeling with ^2^H_3_-serine, averaged as mol% per class (± SD, n=4)**

(XLSX)

**Figure S1.**
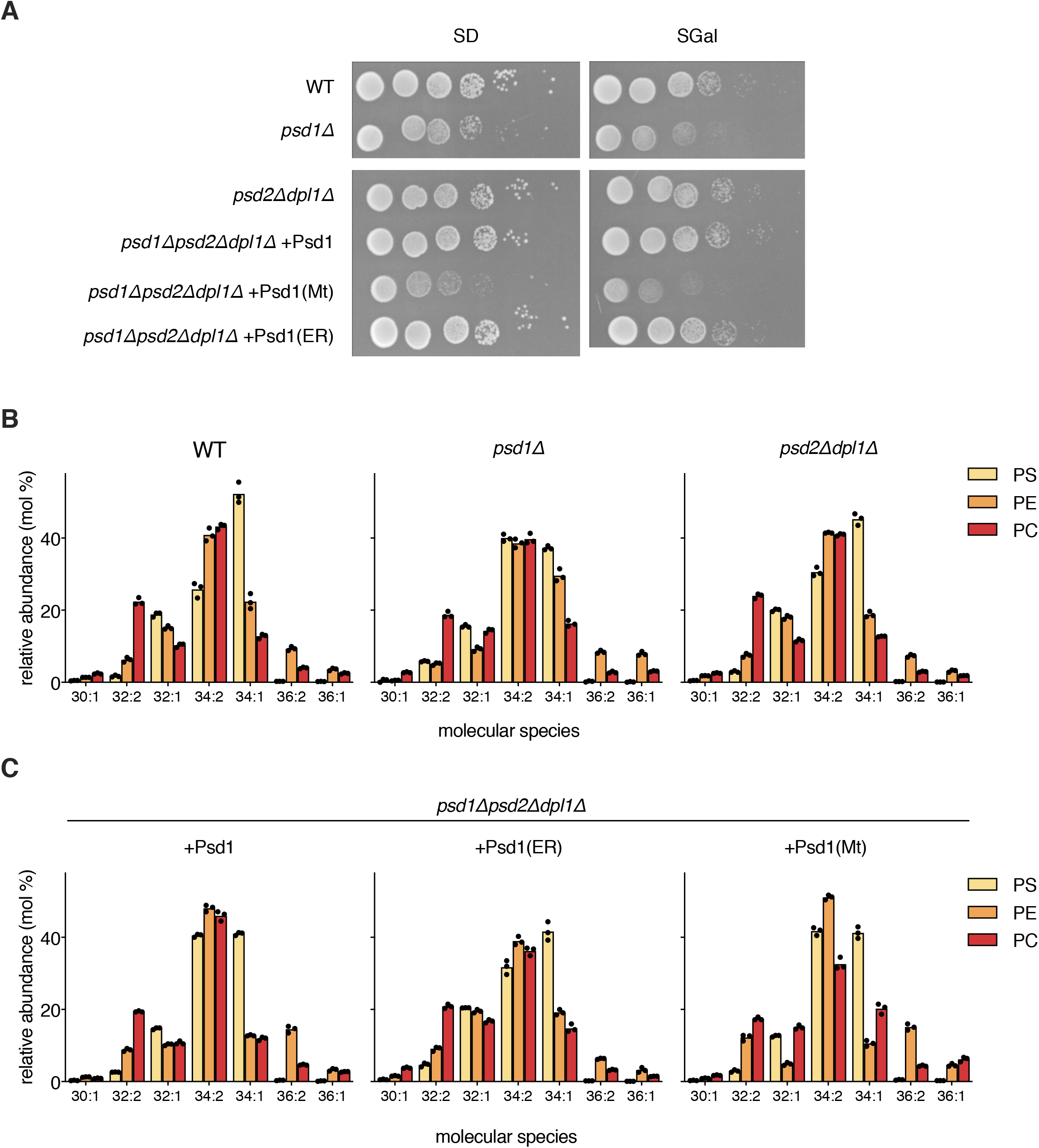
Growth and molecular species profiles of PS, PE and PC of yeast strains defective in PE biosynthesis. (A) Serial dilutions (10^0^ – 10^−5^) of wild type and indicated mutant strains in W303 background were spotted on SD and SGal and incubated at 30°C for 3 days. (B, C) Lipid extracts from the indicated strains prepared after the 20 min pulse (see Figure 2) were analyzed by shotgun lipidomics. Percentages of the molecular species representing at least 2% of PS, PE, or PC are shown. The relatively high C36 content in PE is attributed to contribution of the isobaric phosphatidyldimethylethanolamine species. Data are presented as mean of 3 biological replicates, with the individual values indicated. Numerical data can be found in Data S1.

**Figure S2.**
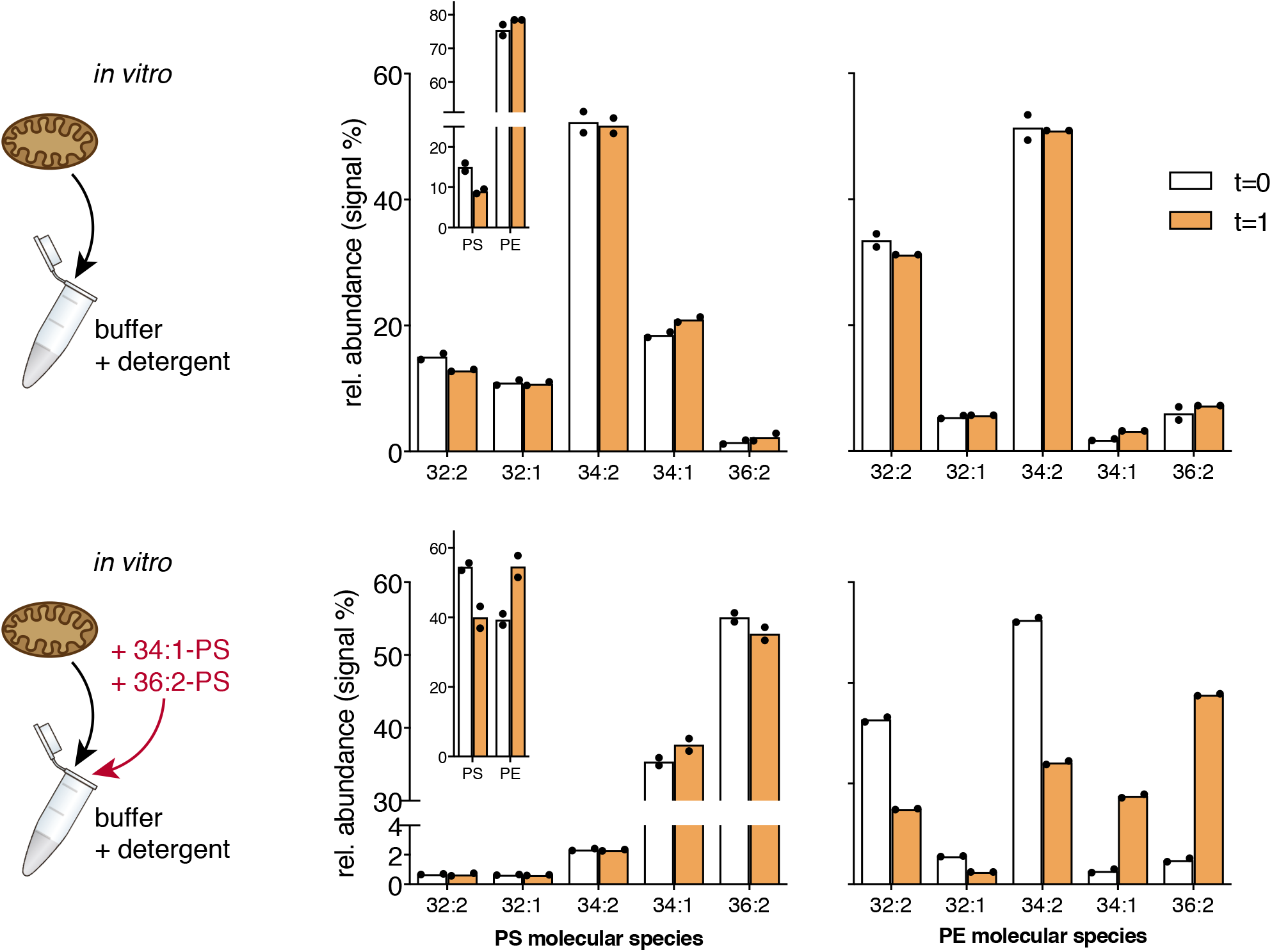
Non-selective conversion of PS molecular species by Psd1 *in vitro*. Molecular species profiles of PS and PE were recorded by LC-MS before and after 1 h of incubating detergent-solubilized mitochondria from a *psd2Δ* strain at 30°C without (top) or with (bottom) the exogenous PS molecular species indicated. The insets comparing the summed signal of PS and PE species as percentage of the total (PS+PE+PI) signal at t=0 and t=1 h, show net conversion of PE to PE. Bars represent the mean of two biological replicates with the individual values indicated. Relative abundance is shown for molecular species that contribute at least 1% of total PS or PE.

**Figure S3.**
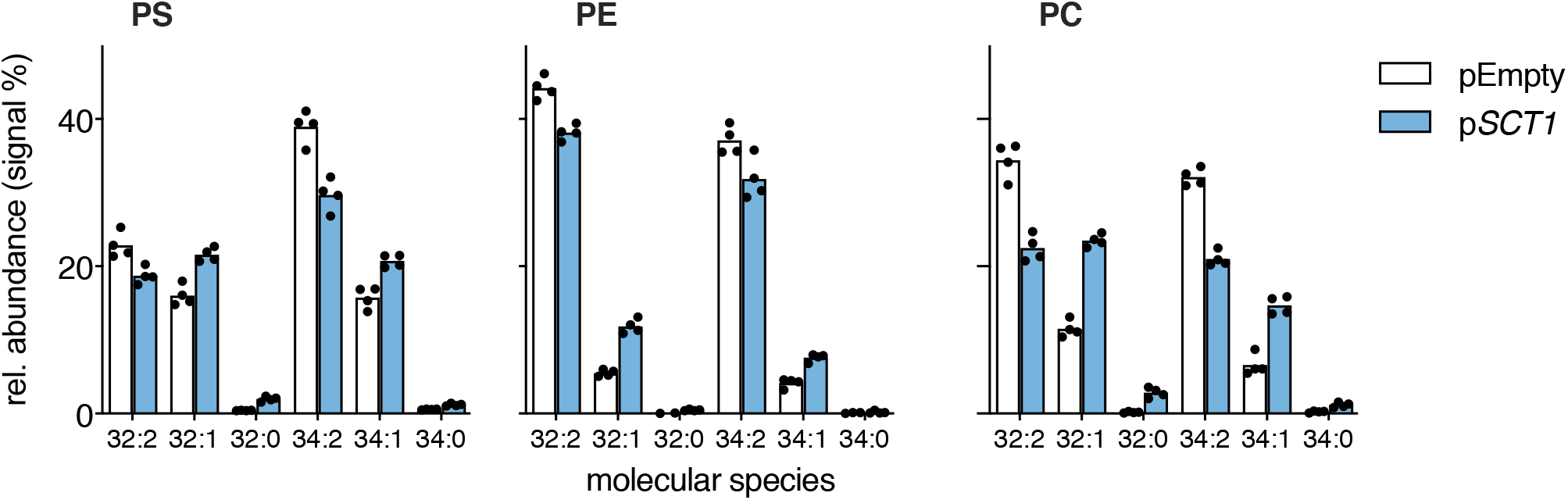
Profiles of the C32 and C34 molecular species of PS, PE, and PC in wild type BY4741 under conditions of *SCT1*-overexpression *versus* control (pEmpty) as determined by ESI-MS/MS. Bars represent the mean of four biological replicates with the individual values indicated. Numerical data can be found in Data S2.

**Figure S4.**
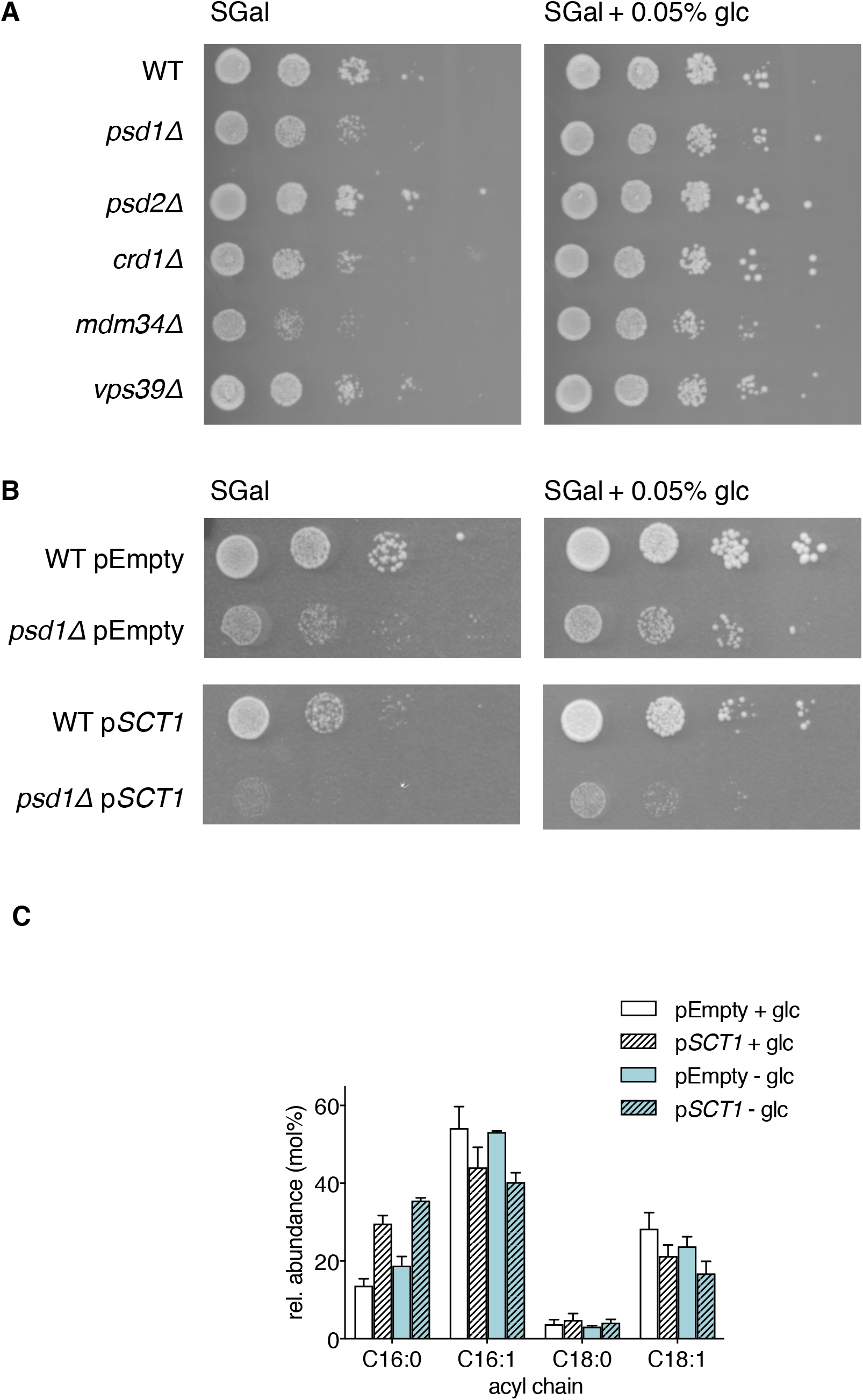
Growth of mitochondrial mutants on synthetic galactose medium is improved by addition of 0.05% glucose. (A) Growth of WT, *psd2D*, and mutant strains disturbed in mitochondrial biogenesis on SGal (left) or SGal + 0.05% glucose (right). Serial dilutions (10^−1^ – 10^−5^) were spotted and incubated for 3 days at 30°C. (B) Growth of WT and *psd1D* overexpressing *SCT1* (p*SCT1*) *vs*. empty vector control (pEmpty) on SGal (left) or SGal + 0.05% glucose (right). Serial dilutions (10^−1^ – 10^−4^) were spotted and incubated for 3 days at 30°C. (C) Acyl chain composition of WT overexpressing *SCT1* (p*SCT1*; dashed bars) *vs*. empty vector control (pEmpty; open bars) cultured in SGal + 0.05% glucose (white bars, data from Fig 3B) or SGal (green bars). Data is depicted as mean + SD (n ≥ 3).

**Figure S5.**
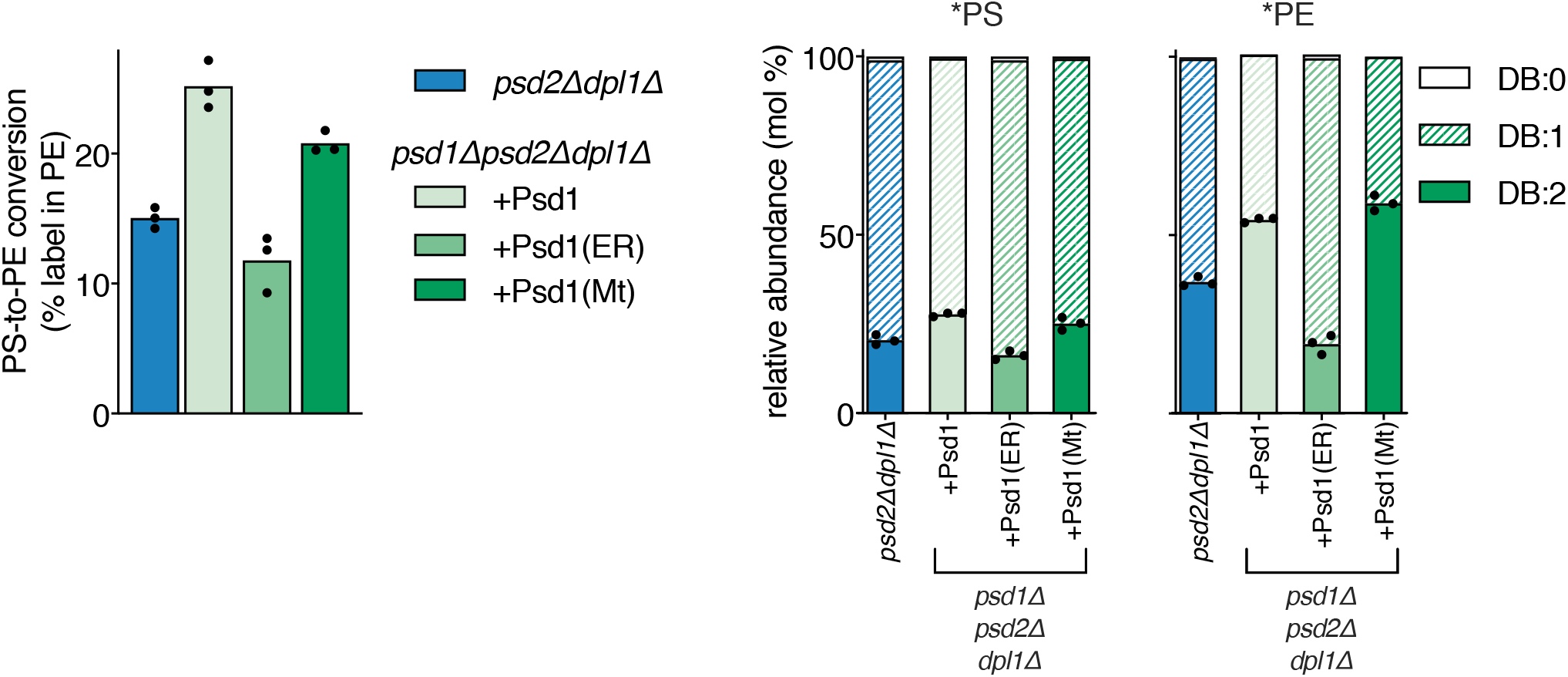
Proportion of di-unsaturated molecular species in newly synthesized PS and PE increases with Psd1 activity in mitochondria. PS decarboxylase activity (data taken from Fig 2A and 2C) and proportions of di-unsaturated (DB:2), monounsaturated (DB:1) and di-saturated (DB:0) molecular species in ^13^C_3_^15^N -labeled PS (*PS) and ^13^C_2_^15^N -labeled PE (*PE) after 20 min incubation with ^13^C_3_^15^N-serine, of the W303 mutant strains indicated. Individual data (n=3) are derived from the experiment shown in Figure 2. Underlying data can be found in Data S1.

**Figure S6.**
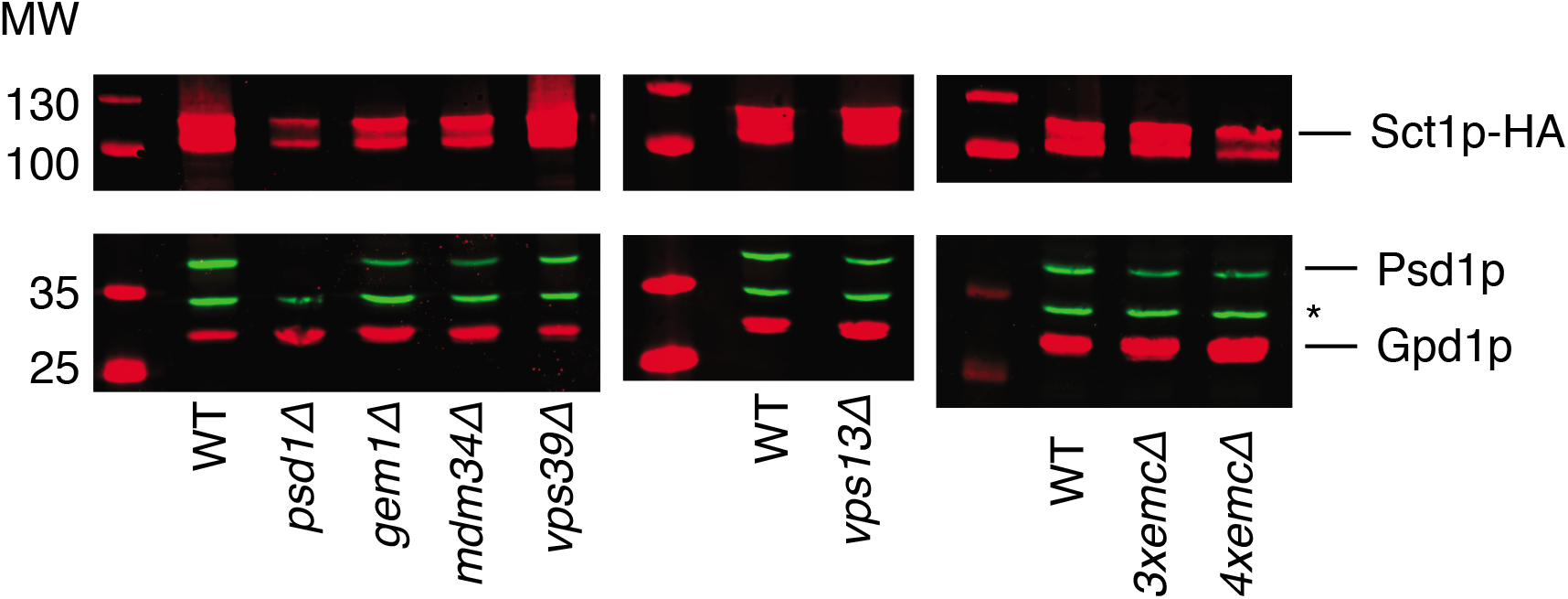
Expression levels of Psd1p and Sct1p in ERMES, vCLAMP and EMCmutants overexpressing *SCT1*. Western blot analysis of the (over)expression of Sct1p and Psd1p in the indicated strains overexpressing *SCT1*. Antibodies used were mouse anti-HA (Sct1p-HA; red), rabbit anti-Psd1p (green), and mouse anti-Gpd1p (loading control; red). * indicates an aspecific staining. Molecular weights (kDa) of the marker bands (Pageruler Prestained, Thermo Scientific) are indicated.

**Figure S7.**
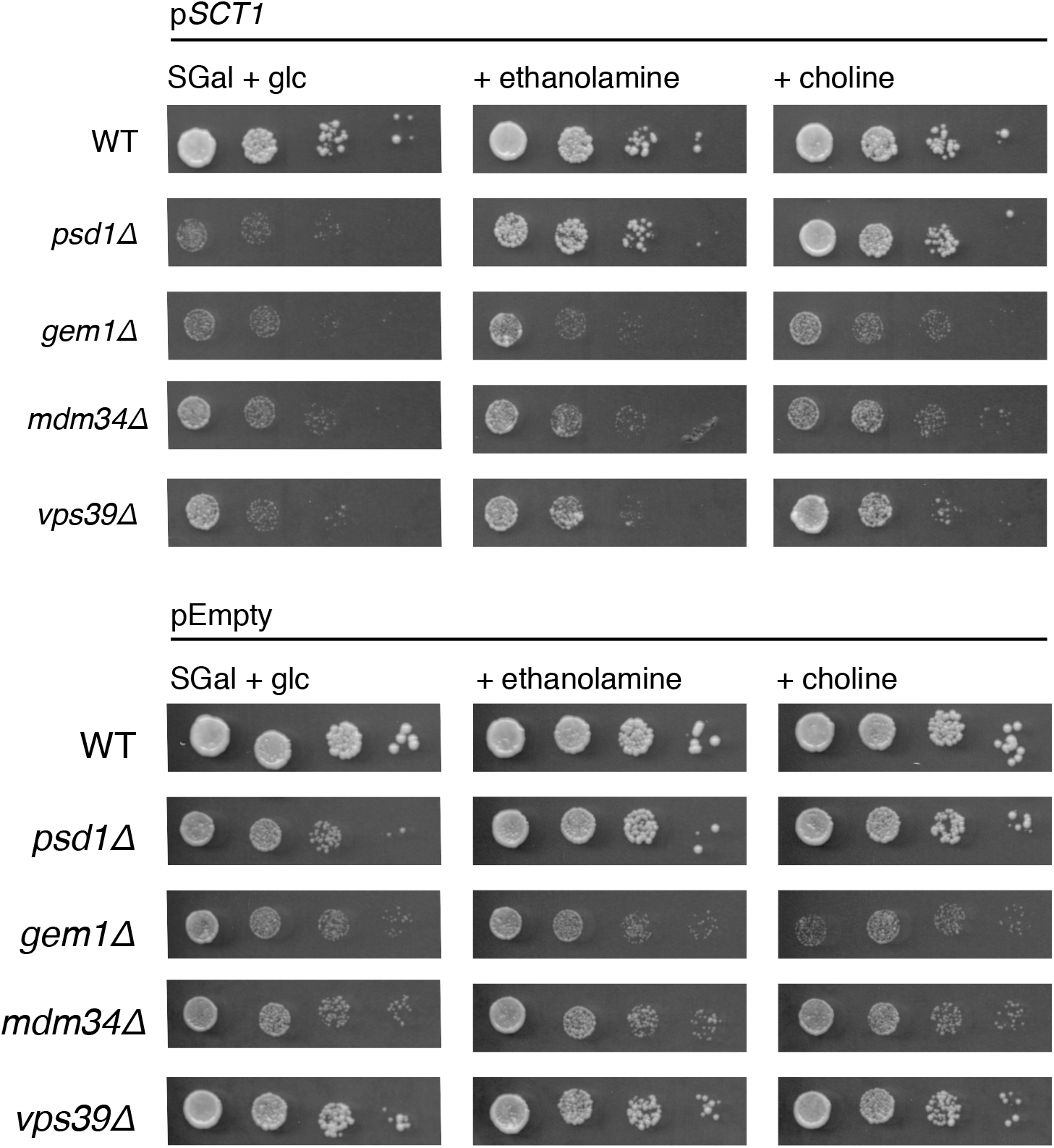
Effect of ethanolamine and choline on the growth of WT and indicated mutants overexpressing *SCT1* (p*SCT1*) *vs*. control (pEmpty). Serial dilutions (10^−1^ – 10^−4^) of the indicated strains were spotted on SGal + 0.05% glucose and incubated for 3 days at 30°C. Ethanolamine or choline (final concentration of 1 mM) were supplied as indicated.

