## Supplementary material for "Molecular species selectivity of lipid transport creates a mitochondrial sink for di-unsaturated phospholipids": Table S1

**Table S1. Acyl chain composition of WT and indicated mutants under *SCT1*-overexpression (p*SCT1*) *vs.* empty vector control (pEmpty)**

All strains were cultured on SGal + 0.05% glucose in the absence or presence of 1 mM ethanolamine ($E^+$) or 1 mM choline ($C^+$). Data is presented as mean (± SD) from n analyses.

| | % C16:0 | % C16:1 | % C18:0 | % C18:1 | SFA / UFA | *n* |
|---|---|---|---|---|---|---|
| **SGal +Glc** | | | | | | |
| WT pEmpty | 13.7 (±1.8) | 54.2 (±5.5) | 3.8 (±1.1) | 28.3 (±4.1) | 0.21 (±0.03) | *15* |
| WT p*SCT1* | 29.3 (±1.8) | 45.3 (±3.6) | 4.5 (±1.0) | 20.9 (±2.6) | 0.51 (±0.04) | *10* |
| *psd1Δ* pEmpty | 18.1 (±2.0) | 51.0 (±5.8) | 5.9 (±1.9) | 25.1 (±3.2) | 0.32 (±0.05) | *15* |
| *psd1Δ* p*SCT1* | 40.2 (±4.8) | 30.7 (±5.3) | 10.1 (±4.2) | 19.1 (±4.6) | 1.02 (±0.13) | *17* |
| *crd1Δ* pEmpty | 15.4 (±0.5) | 49.0 (±7.4) | 4.8 (±1.3) | 30.9 (±5.7) | 0.25 (±0.03) | *4* |
| *crd1Δ* p*SCT1* | 31.1 (±1.8) | 32.8 (±6.5) | 8.9 (±4.2) | 27.2 (±4.2) | 0.67 (±0.08) | *4* |
| *psd2Δ* pEmpty | 14.1 (±0.7) | 50.4 (±5.9) | 4.5 (±1.3) | 30.8 (±3.9) | 0.23 (±0.03) | *3* |
| *psd2Δ* p*SCT1* | 28.7 (±0.6) | 42.3 (±1.6) | 5.6 (±0.7) | 23.4 (±1.4) | 0.51 (±0.01) | *3* |
| *gem1Δ* pEmpty | 13.4 (±1.0) | 50.5 (±6.2) | 4.8 (±1.6) | 31.3 (±5.3) | 0.22 (±0.03) | *7* |
| *gem1Δ* p*SCT1* | 38.7 (±3.4) | 32.0 (±3.2) | 8.7 (±1.8) | 20.5 (±3.3) | 0.91 (±0.12) | *6* |
| *mmm1Δ* pEmpty | 15.2 (±0.8) | 50.2 (±1.3) | 5.8 (±0.6) | 28.8 (±0.8) | 0.27 (±0.02) | *3* |
| *mmm1Δ* p*SCT1* | 35.5 (±2.5) | 33.3 (±8.9) | 9.9 (±4.9) | 21.2 (±4.9) | 0.84 (±0.16) | *6* |
| *mdm10Δ* pEmpty | 13.5 (±1.4) | 50.6 (±10.3) | 5.2 (±3.2) | 30.8 (5.8) | 0.23 (±0.07) | *3* |
| *mdm10Δ* p*SCT1* | 35.0 (±6.0) | 36.5 (±9.8) | 8.1 (±3.1) | 20.5 (±3.2) | 0.79 (±0.25) | *6* |
| *mdm12Δ* pEmpty | 16.3 (±2.2) | 54.8 (±6.6) | 5.2 (±1.2) | 27.4 (±3.8) | 0.22 (±0.05) | *4* |
| *mdm12Δ* p*SCT1* | 34.9 (±2.1) | 32.4 (±3.0) | 9.3 (±1.0) | 23.3 (±3.0) | 0.80 (±0.10) | *4* |
| *mdm34Δ* pEmpty | 14.1 (±1.8) | 49.9 (±5.3) | 5.4 (±1.8) | 30.7 (±3.4) | 0.24 (±0.05) | *7* |
| *mdm34Δ* p*SCT1* | 35.7 (±2.4) | 37.6 (±3.3) | 7.3 (±1.2) | 19.4 (±1.8) | 0.76 (±0.09) | *7* |
| *vps13Δ* pEmpty | 14.3 (±0.9) | 53.6 (±5.1) | 3.4 (±1.0) | 28.7 (±3.3) | 0.22 (±0.03) | *3* |
| *vps13Δ* p*SCT1* | 29.5 (±1.0) | 40.7 (±1.6) | 6.0 (±0.5) | 23.8 (±0.5) | 0.55 (±0.04) | *3* |
| *vps39Δ* pEmpty | 15.3 (±1.1) | 54.6 (±5.8) | 5.5 (±1.5) | 24.6 (±3.8) | 0.26 (±0.04) | *8* |
| *vps39Δ* p*SCT1* | 33.3 (±3.5) | 39.3 (±7.1) | 7.5 (±2.9) | 19.9 (±3.8) | 0.70 (±0.12) | *8* |
| *ypt7Δ* pEmpty | 13.5 (±0.9) | 48.3 (±2.6) | 5.5 (±0.4) | 32.8 (±1.4) | 0.23 (±0.02) | *3* |
| *ypt7Δ* p*SCT1* | 32.5 (±3.4) | 37.7 (±3.0) | 7.2 (±1.5) | 22.6 (±3.8) | 0.66 (±0.07) | *3* |
| *3xemcΔ* pEmpty | 17.9 (±1.8) | 45.0 (±4.2) | 5.3 (±0.9) | 31.8 (±2.4) | 0.30 (±0.04) | *4* |
| *3xemcΔ* p*SCT1* | 23.8 (±1.0) | 43.4 (±3.6) | 5.3 (±0.9) | 27.7 (±2.4) | 0.41 (±0.04) | *4* |
| *4xemcΔ* pEmpty | 23.5 (±2.9) | 43.9 (±7.1) | 6.7 (±2.5) | 25.9 (±1.8) | 0.44 (±0.11) | *5* |
| *4xemcΔ* p*SCT1* | 29.2 (±4.2) | 42.1 (±3.9) | 6.0 (±1.3) | 22.7 (±1.7) | 0.55 (±0.12) | *5* |
| **SGal +Glc $E^+$** | | | | | | |
| WT pEmpty | 14.5 (±1.3) | 52.8 (±6.2) | 4.3 (±1.4) | 28.4 (±4.3) | 0.23 (±0.03) | *3* |
| WT p*SCT1* | 27.8 (±3.5) | 42.2 (±6.3) | 5.3 (±0.8) | 24.8 (±6.1) | 0.50 (±0.07) | *4* |
| *psd1Δ* pEmpty | 15.1 (±1.2) | 53.8 (±2.6) | 3.3 (±0.5) | 27.8 (±3.1) | 0.23 (±0.02) | *7* |
| *psd1Δ* p*SCT1* | 33.1 (±2.7) | 39.5 (±2.2) | 5.2 (±0.8) | 22.3 (±2.5) | 0.62 (±0.08) | *7* |
| *gem1Δ* pEmpty | 16.2 (±3.1) | 48.8 (±3.7) | 5.3 (±1.5) | 29.7 (±5.3) | 0.28 (±0.07) | *3* |
| *gem1Δ* p*SCT1* | 36.7 (±3.2) | 37.7 (±3.4) | 6.2 (±1.1) | 19.4 (±2.1) | 0.76 (±0.11) | *6* |
| *mdm34Δ* pEmpty | 14.9 (±3.6) | 46.6 (±4.5) | 6.3 (±2.0) | 32.2 (±2.5) | 0.27 (±0.09) | *4* |
| *mdm34Δ* p*SCT1* | 31.7 (±5.3) | 34.8 (±9.6) | 8.8 (±5.0) | 24.8 (±9.6) | 0.68 (±0.04) | *3* |
| *vps39Δ* pEmpty | 14.2 (±1.0) | 56.2 (±3.8) | 5.2 (±0.5) | 24.4 (±3.6) | 0.24 (±0.02) | *5* |

| | | | | | | |
|---|---|---|---|---|---|---|
| *vps39Δ* p*SCT1* | 32.9 (±4.1) | 41.3 (±4.9) | 6.4 (±1.7) | 19.5 (±3.8) | 0.65 (±0.11) | *5* |
| **SGal +Glc C⁺** | | | | | | |
| WT pEmpty | 14.1 (±1.8) | 52.6 (±5.8) | 3.1 (±1.1) | 30.3 (±4.6) | 0.21 (±0.04) | *4* |
| WT p*SCT1* | 25.6 (±1.2) | 42.1 (±3.7) | 4.0 (±0.8) | 28.3 (±3.8) | 0.41 (±0.02) | *5* |
| *psd1Δ* pEmpty | 15.0 (±1.8) | 49.7 (±6.4) | 3.8 (±1.1) | 31.5 (±4.6) | 0.23 (±0.04) | *6* |
| *psd1Δ* p*SCT1* | 27.5 (±3.0) | 38.8 (±4.0) | 5.1 (±2.2) | 28.7 (±4.2) | 0.48 (±0.03) | *7* |
| *gem1Δ* pEmpty | 13.9 (±1.0) | 46.7 (±3.4) | 4.2 (±0.5) | 35.2 (±3.8) | 0.22 (±0.01) | *3* |
| *gem1Δ* p*SCT1* | 31.6 (±1.8) | 37.5 (±2.0) | 5.6 (±0.6) | 25.4 (±1.3) | 0.59 (±0.05) | *4* |
| *mdm34Δ* pEmpty | 12.3 (±1.1) | 43.6 (±5.5) | 5.5 (±1.5) | 38.5 (±5.2) | 0.22 (±0.01) | *4* |
| *mdm34Δ* p*SCT1* | 30.3 (±3.0) | 35.7 (±1.2) | 6.5 (±0.3) | 27.5 (±3.8) | 0.58 (±0.07) | *3* |
| *vps39Δ* pEmpty | 15.2 (±2.1) | 50.1 (±6.8) | 6.3 (±1.9) | 28.5 (±4.6) | 0.28 (±0.06) | *5* |
| *vps39Δ* p*SCT1* | 24.2 (±1.5) | 40.8 (±11.1) | 6.9 (±2.9) | 28.1 (±7.2) | 0.46 (±0.08) | *5* |
