## Supplementary material for "Molecular species selectivity of lipid transport creates a mitochondrial sink for di-unsaturated phospholipids": Table S2

**Table S2. Yeast strains and plasmids**

| Strain | Genotype | Source/Reference |
|---|---|---|
| Wild type W303 (JNY30) | MAT**a** *leu2-3,112 trp1-1 can1-100 ura3-1 ade2-1 his3-11,15* | [1] |
| W303 *psd1Δ* (JNY1965) | W303 *psd1::NatMX6* | [1] |
| W303 *psd2Δdpl1Δ* (MA97) | W303 *dpl1::hphNT1 psd2::KanMX6* | [2] |
| W303 *psd1Δpsd2Δdpl1Δ* +Psd1 (JNY1979) | W303 *psd1*::HIS3 *psd2::KanMX6 dpl1::hphNT1 ura3-1::*Psd1 | [1] |
| W303 *psd1Δpsd2Δdpl1Δ* +Psd1(Mt) (JNY1982) | W303 *psd1*::HIS3 *psd2::KanMX6 dpl1::hphNT1 ura3-1::*$Psd1_{MT}$ | [1] |
| W303 *psd1Δpsd2Δdpl1Δ* +Psd1(ER) (JNY1984) | W303 *psd1*::HIS3 *psd2::KanMX6 dpl1::hphNT1 ura3-1::*$Psd1_{ER}$ | [1] |
| Wild type BY4741 | MAT**a** *his3Δ1 leu2Δ0 met15Δ0 ura3Δ0* | EuroSCARF |
| *psd1Δ* | BY4741 *psd1::KanMX* | EuroSCARF |
| *psd2Δ* | BY4741 *psd2::KanMX* | EuroSCARF |
| *crd1Δ* | BY4741 *crd1::KanMX* | EuroSCARF |
| *gem1Δ* | BY4741 *gem1::KanMX* | EuroSCARF |
| *mmm1Δ* | BY4741 *mmm1::KanMX* | EuroSCARF |
| *mdm10Δ* | BY4741 *mdm10::KanMX* | EuroSCARF |
| *mdm12Δ* | BY4741 *mdm12::KanMX* | EuroSCARF |
| *mdm34Δ* | BY4741 *mdm34::KanMX* | EuroSCARF |
| *vps39Δ* | BY4741 *vps39::KanMX* | EuroSCARF |
| *vps13Δ* | BY4741 *vps13::KanMX* | EuroSCARF |
| *ypt7Δ* | BY4741 *ypt7::KanMX* | EuroSCARF |
| *3xemcΔ* (YSL6) | BY4741 *emc1::HIS5 emc2::hygMX4 emc5::kanMX4* | [3] |
| *4xemcΔ* (YSL27) | BY4741 *emc1::HIS5 emc2::hygMX4 emc3::his emc6::kanMX4* | [3] |
| **Plasmid** | | **Source/Reference** |
| pYES2 (pEmpty) | | Invitrogen |
| pYES2-*SCT1*-HH (p*SCT1*) | | [4] |
| pYES2-*PSD1*-HA | | [5] |
| pYPGK18 | | [6] |
| pYPGK18-*PSD1-HA* | | This study |
